# Natural Circulation of Tick-borne Severe Fever with Thrombocytopenia Syndrome Virus in the City Ecosystem, China

**DOI:** 10.1101/2023.07.02.547301

**Authors:** Xing Zhang, Chaoyue Zhao, Xiaoxi Si, Qiang Hu, Aihua Zheng

## Abstract

Severe fever with thrombocytopenia syndrome virus (SFTSV) is rapidly expanding its range in China, because of the accelerated spread of the parthenogenetic *Haemaphysalis longicornis*, Asian long-horned tick (ALT). In this letter, we report the urban circulation of SFTSV between ALTs and hedgehogs in Beijing, China.

**Highlights:** Hedgehogs and ALTs can maintain the natural circulation of SFTSV in the city ecosystem.

Hedgehogs and ALTs are becoming common in Beijing.

Parthenogenetic ALTs are discovered in Beijing.

## Introduction

Severe fever with thrombocytopenia syndrome virus (SFTSV) is a newly emerging tick-borne Bandavirus that can cause severe symptoms in patients include fever, gastrointestinal symptoms, thrombocytopenia, leukopenia and even death [1]. SFTSV was first identified in China in 2009, and has subsequently been found in South Korea [2], Japan [3], Vietnam [4], Myanmar [5] and Pakistan [6] in Asia. Initial outbreaks of SFTSV were mostly restricted to people from remote and mountainous country areas, with people from cities being relatively unaffected [7]. This was certainly true for Beijing residents, where no cases of Severe fever with thrombocytopenia syndrome (SFTS) were reported before 2021.

The Asian long-horned tick (ALT), *Haemaphysalis longicornis*, is the primary vector for SFTSV and the dominant human-biting tick in the SFTSV-endemic areas [8]. ALTs have both bisexual and parthenogenetic populations among which the parthenogenetic populations are widely distributed in China and strongly correlated with the distribution of SFTS cases [9]. Our results suggest that the parthenogenetic ALTs, are probably transported to naïve areas by migratory birds, and this mechanism plays a major role in the long-range spread of SFTSV [9]. Even though many domestic and companion animals show high SFTSV seroprevalence, our studies strongly suggest that only hedgehogs are the major amplifying host for SFTSV in China [10]. Hedgehogs are widely distributed throughout China, encompassing areas where SFTSV is endemic. Due to fast urbanization and strict ecological conservation, many cities, including Beijing, have recently experienced a rapid increase in hedgehog and ALT populations (10, 11). This is a major concern since it implies that SFTSV can now circulate in urban as well as rural areas.

## Result

In October 2021, a suspected case of SFTS was reported at Beijing You’an Hospital, whose samples were tested positive for SFTSV RNA and antibody by the Beijing Center for Disease Control and Prevention [11]. A case report was published in Chinese in *International Journal of Virology* but due to the COVID-19 pandemic was likely overlooked. The 69-year-old male patient lives in Longquanwu Village,Mentougou District, Beijing, where is only 20 miles away from the center of Beijing City and is fully urbanized, with no poultry and livestock present (Fig. 1A).

**Figure 1.**
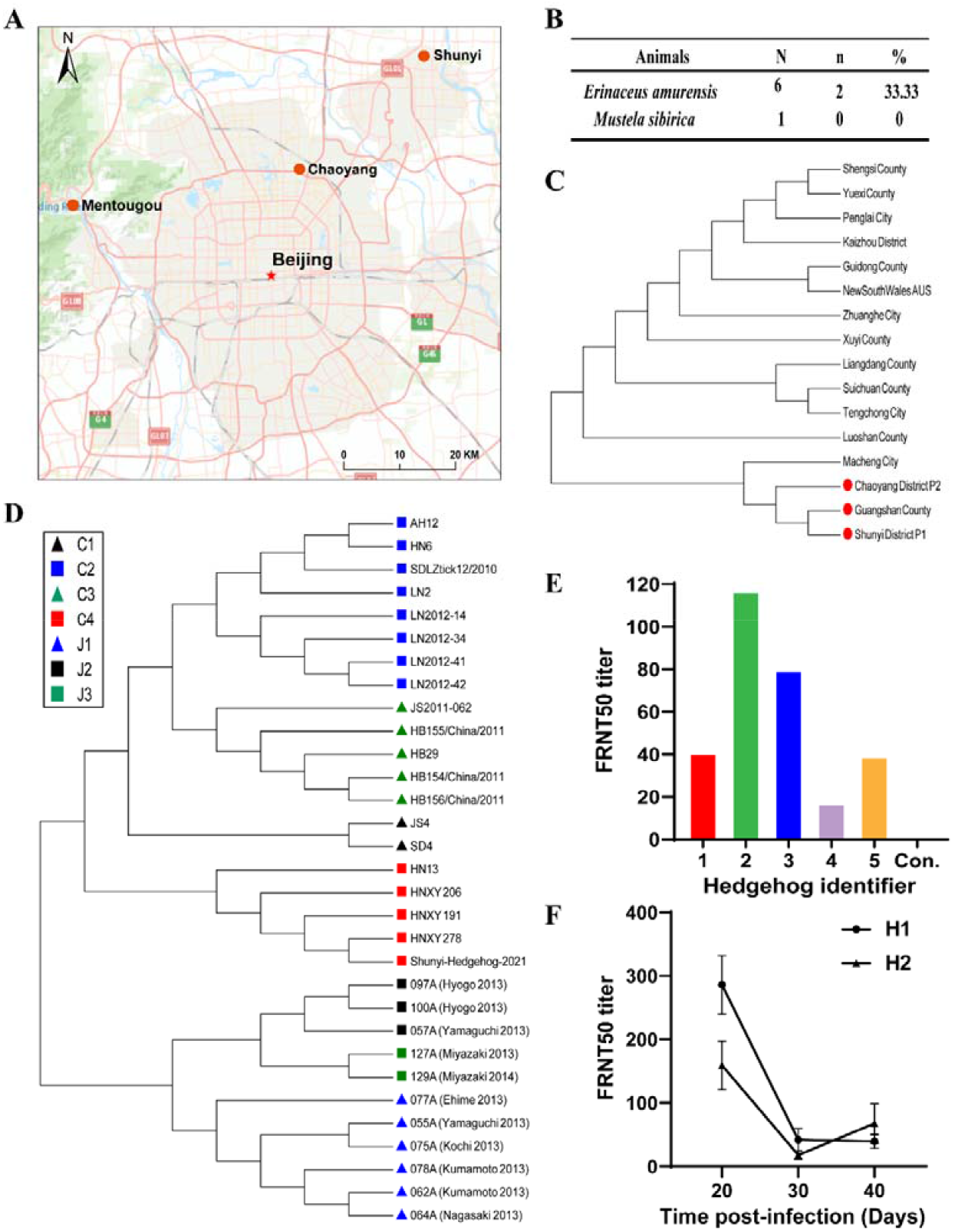
Natural circulation of SFTSV in Beijing City. (A) Sample collection sites of small park in Shunyi District (Shunyi), the Olympic Forest Park in Chaoyang District (Chaoyang), Longquanwu Villiage in Mentougou District (Mentougou). (B) Seroprevalence of animals caught in Shunyi district. N, number of sampled animals; n, number of sampled animals positive for SFTSV antibody; %, percentage of sampled animals positive for SFTSV antibody. (C) Phylogenetic analysis of the parthenogenetic population of ALTs. Maximum likelihood tree was established with the mitochondrial genomes of ALTs collected in Chaoyang District and Shunyi District (Chaoyang District P2, accession number, OL335942 and Shunyi District P1 accession number, OL335941) (red) and from SFTS endemic areas [9]. (D) Maximum likelihood tree was established with the L Segments of SFTSV isolate in Shunyi (Shunyi-Hedgehog-2021) and isolates from SFTS endemic areas [9]. HNXY206, HNXY191, and HNXY278 were isolated from Xinyang City, Henan Province. SFTSV lineages were illustrated by colors and shapes. (E) Neutralizing antibody titer (FRNT50) against SFTSV Wuhan strain in *E. amurensis* hedgehogs caught near Longquanwu Village. Control (Con.) is naïve *E. amurensis* hedgehogs maintained in the lab. The detection limit is 10. (F) Two *E. amurensis* Hedgehogs (H1 and H2) were intraperitoneally challenged with 4 × 10^6^ FFU of SFTSV Wuhan strain and SFTSV neutralizing antibody titer (FRNT50) were monitored at 20, 30, 40 days post infection.

In 2021, two field surveys were conducted to investigate the small mammals and parasitic ticks at two locations in Beijing City. The first location was a small park in Shunyi District, where is surrounded by up-market gated communities, and the second location was the Olympic Forest Park in Chaoyang District, where parthenogenetic ALTs had previously been recorded in 2019 (Fig. 1A) [9]. The Shunyi park is quite small, about 0.5 km^2^, and has a relatively simple ecosystem. The small mammal survey showed only *Erinaceus amurensis* hedgehogs and *Mustela sibirica* yellow weasel in this park (Fig. 1B). On October 19^th^, 2021, six *E. amurensis* hedgehogs were caught in the Shunyi park, two of which (33%) were seropositive for SFTSV (Fig.1B). In addition, ALTs collected from these hedgehogs were positive for SFTSV RNA and an L segment was sequenced (GenBank accession number: OL518989). Sequencing showed that the Shunyi SFTSV (Shunyi-Hedgehog-2021) was clustered into lineage C4, which is similar to the strain of SFTSV collected in Xinyang City, Henan Province, one of the original SFTS hot spots (Fig.1C). In contrast, no SFSTV RNA or antibody was detected in hedgehog serum samples or parasitic ticks collected at the Chaoyang sample site.

Phylogenetic analysis of the whole mitochondrial sequences from the parthenogenetic ALTs collected at Shunyi (Shunyi District P1) and Chaoyang (Chaoyang District P2), further established that these ticks were closely related to those from Guangshan County, Xinyang City in Henan Province (Fig. 1D). The results of the viral sequencing and phylogenetic analysis, when taken together, strongly suggest that both the tick and SFTSV collected in Shunyi District originated from Xinyang City, where the first SFTS case in China was reported. Although no local SFTS cases had been reported in Beijing at that time, our results suggested that a population of hedgehogs and ALTs existed in the Shunyi District that could maintain SFTSV in an urban ecosystem.

In April 2023, a field survey was conducted in Longquanwu Villiage, Mentougou District, around the 2021 patient’s house [11]. Five *E. amurensis* hedgehogs were caught, which showed 100% SFTSV seroprevalence with SFTSV neutralizing antibody titers (FRNT50) ranging from 20 to 110 (Fig. 1E). In addition, 190 ALTs were collected from vegetation and the hedgehogs, all of which was negative for SFTSV RNA (Supplementary Table 1). Interestingly, the sequencing results of mitochondrial DNAs from 40 ALTs showed that they were all bi-sexual.

The immune response of hedgehogs against SFTSV is unusual. In contrast to the stable humoral immune response of experimental dogs [12], the neutralizing antibody titer of *E. amurensis* hedgehogs decreases quickly and almost eliminated by day 40 after intraperitoneal inoculation with SFTSV (Fig. 1F). Thus, it is plausible that active transmission existed between the hedgehogs and ALTs around the patient’s location, although no SFTSV RNA positive ALTs were detected.

## Discussion

Our previous work raised two hypotheses, the first that parthenogenetic ALT are strongly associated with SFTSV disease outbreaks [9] and the second that SFTSV can be maintained between ALTs and hedgehogs in the urban ecosystem in China [10].

In our state-wide survey of ALTs in 2019, parthenogenetic ALTs were quite uncommon in Beijing, where no SFTS cases were reported at that time. Parthenogenetic ALTs were only found in a small park in Shunyi District, and the Olympic Forest Park in Chaoyang District in Beijing [9]. Two years later, SFTSV was detected in the ALTs collected in the Shunyi park. In our opinion, these parthenogenetic ALTs were probably imported to Beijing from Xinyang City, the hot SFTS endemic area, by migratory birds, since Beijing is located on the East Asian-Australasian Flyway. In 2021, the first SFTS patient diagnosed in Mentougou District, where is not far from the Shunyi location. Considering that the patient had not travelled more than 1km from home and the ALT density was high in his neighbourhood, we speculate that he got infected by ALT bites near his house.

Shunyi park has a quite simple ecosystem where is surrounded by up-market gated communities. Only *E. amurensis* hedgehogs and yellow weasel (*Mustela sibirica*) were trapped there. *E. amurensis* hedgehogs were common in Shunyi District and 33% of the hedgehogs collected from the Shunyi park were seropositive for SFTSV. Since there are almost no livestock, poultry, and stray dogs in this location, our discovery reinforces our hypothesis that SFTSV circulation can be maintained in an urban area by just hedgehogs and ALTs.

Longquanwu Village is close to the mountains, but already urbanized, with no poultry and livestock, and is only 26 miles away from the Shunyi park. Hedgehogs caught near the village showed 100% SFTSV seroprevalence. However, so far, no SFTSV RNA has been detected from both ALTs collected from vegetation or hedgehogs. In experimentally infected hedgehogs, the neutralizing antibody titer decreased quickly after SFTSV infection and almost eliminated by 40 days post infection (Fig. 1F). Thus, we believe that all the hedgehogs were infected by ALT bites in this spring. Maybe, the SFTSV-RNA positive rate is too low in this area and more ticks need to be tested in the future, which may also explain the low human case numbers in Beijing. Another hand, migratory birds also may play an important role in natural circulation of tick-borne Severe Fever with Thrombocytopenia Syndrome Virus in the City Ecosystem of China. The more intensive investigation of seroprevalence and SFTSV RNA in migratory birds also need to be done. Overall, our results suggest that the natural circulation of SFTSV have been established in Beijing and need more surveillance.

## Acknowledgments

We thank the following funders: National Key R&D Program of China (2021YFC2300903, 2022YFC2601603), Open Research Fund Program of the State Key Laboratory of Integrated Pest Management (IPM2215), Guangdong Provincial Key R&D Program, No. 2022B1111040001. We thank Dr. Kevin Lawrence from Massey University, New Zealand for reading and revising the manuscript.

